# The smooth transition from many-legged to bipedal locomotion - Gradual ground force reduction and its impact on total ground reaction forces, body dynamics and gait transitions

**DOI:** 10.1101/2021.07.07.451417

**Authors:** Tom Weihmann

## Abstract

Most terrestrial animals move with a specific number of propulsive legs, which differs between clades. The reasons for these differences are often unknown and rarely queried, despite the underlying mechanisms being indispensable for understanding the evolution of multilegged locomotor systems in the animal kingdom and the development of swiftly moving robots. Moreover, when speeding up, a range of species change their number of propulsive legs. The reasons for this behaviour have proven equally elusive. In animals and robots, the number of propulsive legs also has a decisive impact on the movement dynamics of the centre of mass. Here, I use the leg force interference model to elucidate these issues by introducing gradually declining ground reaction forces in locomotor apparatuses with varying numbers of leg pairs in a first numeric approach dealing with these measures’ impact on locomotion dynamics. The effects caused by the examined changes in ground reaction forces and timing thereof follow a continuum. However, the transition from quadrupedal to a bipedal locomotor system deviates from those between multilegged systems with different numbers of leg pairs. Only in quadrupeds do reduced ground reaction forces beneath one leg pair result in increased reliability of vertical body oscillations and therefore increased energy efficiency and dynamic stability of locomotion.

**Significance statement:** The model grants access to the effects of gradual ground force reduction on total ground reaction forces, body dynamics and gait transitions.

## Introduction

In legged terrestrial animals, as well as legged machines moving on level ground, body dynamics are characterised by the ground reaction forces (GRF) applied by the single legs and the temporal coordination between them (i.e. leg coordination patterns (*1*)). By and large, with low external friction body dynamics are directly equivalent to the sum of all forces applied to the ground, particularly at lower and cursorial speeds (*2*). By using a simple numerical approach to study the interaction of single leg ground forces in polypedal locomotor apparatuses, it has been shown previously that the number of walking legs involved significantly affects the impact of ipsilateral phase shifts onto overall ground force oscillations and body-dynamics accordingly (*3*).

When accelerating from low to medium speeds, many few legged animals (i.e. those with less than or equal to four pairs of walking legs like vertebrates, insects and arachnids) shift ipsilateral phase relations (θ) from intermediate values towards values close to 0.5. Ipsilateral phase values describe the occurrence of a leg’s stance phase within the cycle period of an adjacent leg on the same body side. With θ = 0.5, and similar contralateral phase relations of a pair of pacemaker legs (which is typical for so-called symmetrical gaits like trot and its multilegged equivalents (*4*)), two alternating sets of temporally synchronised sets of legs emerge. However, at medium and higher velocities, many vertebrates, and also a range of insect and arachnid species, shift ipsilateral phase relations (θ) significantly away from strict alternation (θ = 0.5). Deviations from strictly alternating sets of legs result in reduced oscillations of the total vertical forces (Fig. 1) and vertical oscillations of the animals’ bodies accordingly. They are particularly advantageous when leg elasticities cannot be used owing to anatomical constraints or specific environmental conditions (*3, 5-7*).

**Fig. 1.**
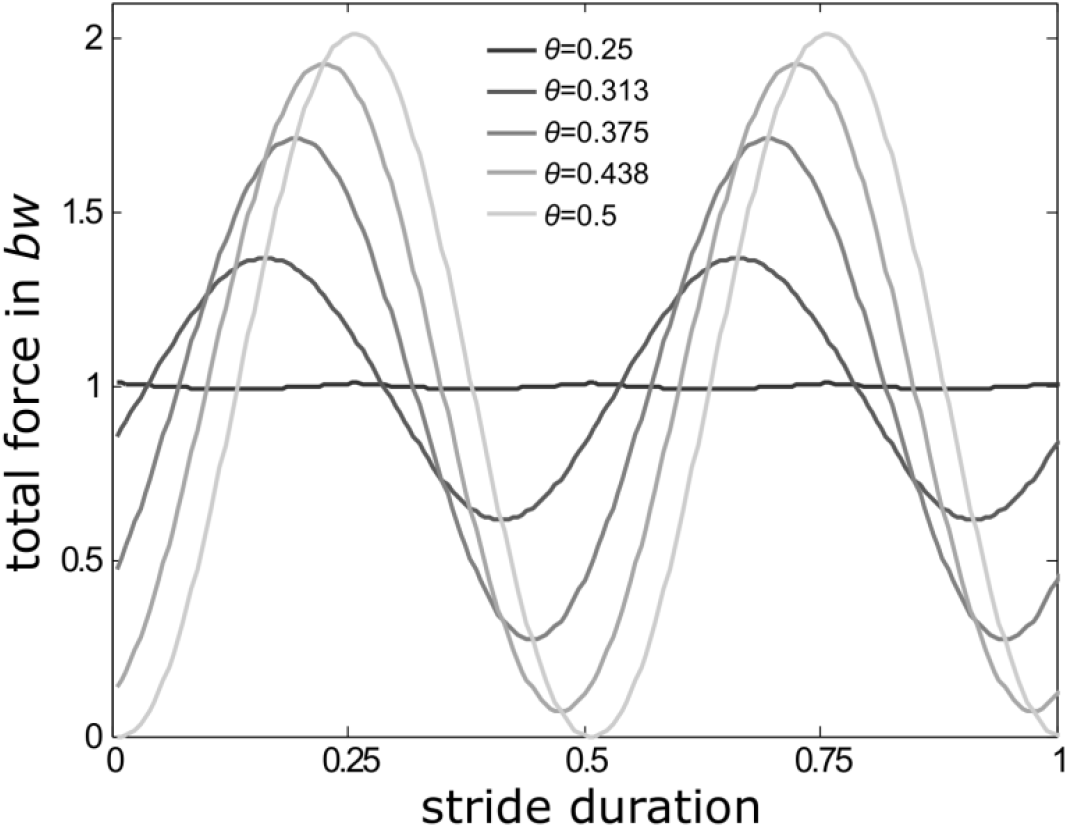
Reduction of total force amplitudes with changing ipsilateral phase relations in a quadrupedal locomotor system moving with stance and swing durations of equal length, i.e. with a duty factor of 0.5. At θ = 0.5 (light grey force trace), i.e. alternating sets of legs (trot), the consecutive sine-shaped GRF-traces coincide with the stance phases of the two successively active sets of legs. The more θ deviates from 0.5, GRFs increasingly blend into each other and total force amplitudes decline to reach a minimum with only minor oscillations around body weight (*bw*) at θ = 0.25 (dark grey line).

Strict alternations of synchronised sets of legs are typical for medium-speed bouncing gaits, like trot or its polypedal equivalents. These gaits provide efficient storage and recovery of movement energy during stance (*8*) and are currently widely employed for faster modes of locomotion in legged robots (e.g. *9, 10*).

In animals, neuronal networks with their enormous intrinsic variability can generate almost arbitrary leg coordination patterns. Evolution has brought forth similar rhythmic patterns of limb like appendages in a number of phylogenetically only loosely related groups, in which some have even evolved their legs independently of each other (e.g. *4, 11, 12-15*). Manmade machines and models mostly adopt coordination patterns that closely mimic those found in animals. Apparently, the interaction with the physical world and the constraints implied by physical laws, therefore, is more important than specifically how leg coordination is achieved (cp. *16*).

Many polypedal animals reduce GRF gradually beneath some (mostly the anterior) of their walking legs, in particular at higher running speeds (*17-19*). In some lineages, differential leg use has led, via transitional forms, to varying numbers of propulsive leg pairs. Thus, in mammals (e.g. men, pangolin) and archosaurs, quadrupedal ancestors have repeatedly brought forth bipedal forms while some lineages of dinosaurs went the reverse direction and regained quadrupedalism (*20*). The evolution of arthropods also seems to be characterised by repeated and independent reductions in the number of propulsive legs. Particularly in small terrestrial forms, these reductions have probably extensive advantages regarding locomotion energetics and control effort, and therefore may have influenced major morphological changes in the course of these animals’ transitions from marine to terrestrial habitats (*21, 22*).

Among terrestrial arthropods, reductions in propulsive contributions of certain pairs of legs can be observed specifically in fast moving insects, crabs and arachnids. For the cockroach *Periplaneta americana*, concurrent increases in running speed and the body’s angle to the substrate have been reported (*17*). An increasingly upright body posture tends to successively reduce the ground contact of the relatively shorter anterior leg pairs, which results in decreasing duty factors (the quotient of stance phase duration and cycle duration of a leg) of these legs and, finally, can even lead to bipedal runs. In the mainly used running legs, lower duty factor values usually refer to higher running speeds and vice versa. Different species of ghost crabs have been observed to reduce the number of propulsive legs with increasing running speeds, which can also result in bipedal runs (*19, 23*); although here the longest legs of the two sides of the body alternate with each other. In arachnids, a couple of lineages exist that typically use only three of their four pairs of legs for locomotion (*1*). Thus, harvestmen use their second legs for propulsion when climbing steep slopes (*24*), while these legs are solely of sensorial nature on shallower substrates (*25*). However, also in spiders that usually run with all four pairs of their legs, stance durations and amplitudes of the legs’ GRFs change with speed resulting in significantly reduced impulse contributions of the forelegs when running at high velocities (*26*). Moreover, carrying loads with anterior or posterior devices or uneven mass distribution in general (*27, 28*), likely affects the loading conditions of the different pairs of legs that are distributed in anterior posterior directions along the body axis of animals or technical implementations accordingly.

Currently, it is largely unclear why animals change the number of propulsive legs when changing running speeds, however, since the behaviour has become established in a wide range of animals there seem to be some associated advantages. Energetically there are two different ways to optimise locomotion: i) using vertical oscillations of the body’s centre of mass (COM) for storage-recovery cycles of kinetic and potential energy (*29*) or ii) keeping COM height as constant as possible in order to avoid losses (*30*). For attaining oscillation dynamics, the synchronisation of functional sets of legs is a requirement that enables efficient repulsion dynamics of the COM against gravity. When body dynamics follow that of an inverted pendulum, the COM reaches its highest position during mid-stance and movement energy is stored as potential energy, whereas the COM reaches its lowest position and movement energy is elastically stored, when body dynamics are similar to that of a spring mass system.

When animals move at constant speeds, vertical components of single leg GRF are typically much larger than fore-aft and lateral forces, making them particularly suitable for use in energy storage and recovery cycles. This is true also in animals with sprawled legs such as reptiles or arthropods like insects and arachnids (*8, 25, 31, 32*). Accordingly and in absolute terms, overall vertical force amplitudes are particularly affected by phase shift induced ground force interferences, which makes them most decisive and characteristic for changes in COM dynamics and therefore gaits (*1, 21*). In many biological and synthetic systems, vertical GRF are approximately symmetrical around midstance (e.g. *1, 8, 23, 29, 31, 33-35*) in particular when duty factors are not too high (but see the discussion section for the impact of deviations). Since ample time is available for the swing phases, in locomotor systems with many leg pairs, low duty factors can occur already at relatively low running speeds (*36*), making symmetrical, one-humped single leg GRF traces the dominant pattern in a wide range of species.

The lower the number of propulsive legs synchronously in contact with the ground and the less sprawled leg positioning is, the larger are the vertical force components compared to lateral and fore-aft force components. However, animals with higher numbers of leg pairs like isopods, centipedes and millipedes, i.e. those who are likely to have a relatively high ratio of horizontal to vertical GRF, often have burrowing life styles and typically inhabit crevices and burrows. Most of them regularly have to deal with much higher counterforces, such as substrate friction (*37-39*), than surface dwellers. Both, required propulsive force generation as well as bearing of body weight, here are distributed among many legs. In line with model predictions (*3*), coordinative changes in leg activity in these animals do not affect vertical, i.e. sagittal plane, body dynamics and regular vertical oscillations of the COM are largely missing.

The force interference model for integer numbers of walking legs (*3*) predicts decreasing total force amplitudes when ipsilateral leg coordination deviates from the strict alternating pattern. Shifting away from perfect alternation, results in symmetric minima of the total force amplitudes (cp. also Fig. 2, left column). The position of these minima in the phase range are specific for the numbers of propulsive legs and closer to ipsilateral phase values of 0.5 the more legs an animal has. These minima are exploited by quadru-, hexa- and octopedal animals for energetic optimisation of their fast locomotion if storage-recovery cycles of kinetic and potential energy are not applicable (*1, 3, 32*).

**Fig. 2.**
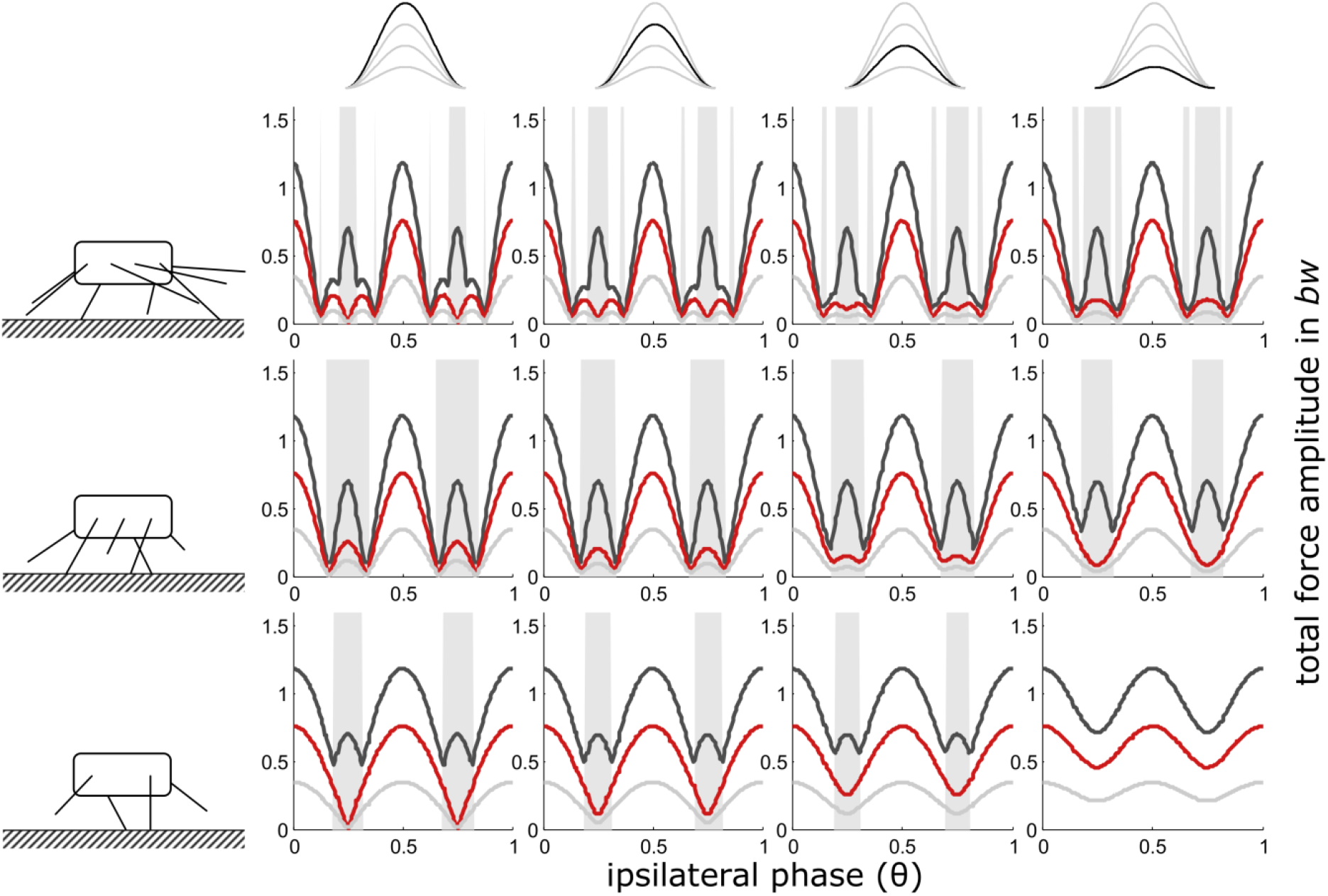
Constant contact durations (i): Dependency of total vertical leg force amplitudes on the phase shift of ipsilateral adjacent legs and duty factor with gradually decreasing contributions of increasing numbers of leg pairs. Duty factors shown: 0.3 (dark grey), 0.5 (red) and 0.8 (light grey). The top row indicates the degree of reduction in the pair of legs with reduced GRF and the type of force alignment (here (i)). Accordingly, in the third column from the right, the impulse beneath one leg pair is reduced by ¼, in the 2^nd^ column from the right by ½ and in the rightmost column by ¾. The pictograms to the left indicate the number of propulsive pairs of legs valid for the respective row of subplots, i.e. four in the upper row, three in the middle row and two in the bottom row. With low duty factors and intermediate phase shifts, the peak frequency of the force oscillations deviates from two times the stride frequency; these intervals are shaded in grey.

The impact of the number of walking legs on body dynamics relies on general physical and physiological laws and principles. One of these principles shall be further explored in the present study. For this purpose, the leg force interference model (*3*) was adapted by implementing gradual reductions in the GRF of single leg pairs in relation to the remaining legs. This model extension allows scrutinising the effects of such reductions on overall body dynamics and examining the mechanisms that govern changes in the number of propulsive legs in the range from truly polypedal animals, like isopods, to quadrupeds that occasionally reduce the forces applied by their forefeet.

Animals that reduce the number of propulsive legs when speeding up can either keep the position of their body axis unaltered (e.g. lizards, *18*) or increase the body’s angle of attack (e.g. cockroaches, *17*). With alternating sets of legs, constant body orientation rather leads to mid-stance synchronisation whereas increasing body angles result in delayed foreleg touch-downs with synchronised take-offs. In other, mostly polypedal, species with uneven force distribution among the legs, rearward directed legs might also take-off prematurely.

Accordingly, four different regimes of force reduction were explored: i) where impulse reduction was achieved by reduction of the force amplitudes alone and ii to iv) where impulse reduction was achieved while maintaining the GRF’s general shape, i.e. a cosine wave derived from previous experimental results and modelling approaches (*8, 34, 40 and see methods*). Shape maintenance and simultaneous impulse reduction resulted in shorter contact durations. The timing of shorter contacts was aligned with either ii) touch down iii) mid stance or iv) take-off of an assumed unchanged step.

With synchronised sets of otherwise alternating sets of legs, the different reference points imply aligned touch-downs, mid-stances or take-offs of the complete set of legs. When ipsilateral phase relations were then changed, the alignment diminished accordingly; nevertheless, the initial alignment has consequences on total force amplitudes for the whole range of leg coordination patterns.

## Results

When all legs generate the full amount of GRF, the results of the present approach cover those presented by Weihmann (*3*); however, when the forces beneath one of the leg pairs diminish, the results can deviate significantly. With constant contact durations, the effects of gradual GRF reduction were largely restricted to quadrupeds. Reduced contact durations, as found when the GRF shape was maintained, resulted in more pronounced deviations from the patterns found when leg numbers changed by integers. Here, too, the deviations increase as the number of legs decreases.

### i) Reduction of force amplitude and constant contact durations

The width of the central ranges of the phase graphs, with the maxima of the total force amplitudes at phase values of 0.5 (strict alternation of synchronized sets of legs), increases steadily from high numbers of pairs of walking legs towards lower numbers (Fig. 2). However, during the transition from two to only one pair of propulsive legs the widening of the central range comes to an end and only the extent of the force amplitude minima decreases. Their positions in the phase range remain constant at θ = 0.25 and θ = 0.75. In quadrupedal locomotion (2 pairs of walking legs), every reduction of GRF in one pair of legs leads to diminishing minima of total force amplitude until, eventually, the force amplitude becomes constant when only 2 legs hit the ground alternatingly. With only one remaining pair of legs, naturally, ipsilateral phase shifts, and therefore the present model, are no longer applicable (see methods section). Here, other model approaches are better suited to investigate the dynamics of the COM (cp. *41*).

In contrast to the force reduction schemes with shape consistency (see below), the maxima of total force amplitudes for the different numbers of walking legs remained constant for a given duty factor when the contact durations of all leg pairs were constant.

As with integer numbers of pairs of legs (*3*), oscillation frequencies of more than twice the stride frequency occur predominantly at low duty factors (0.3) at intermediate ipsilateral phase relations (Fig. 2). Affected phase values range between the amplitude minima of total forces that occur just adjacent to the force amplitude maxima at leg synchrony (θ = 0 or 1) and leg alternation (θ = 0.5). With less than four fully contributing legs, high oscillation frequencies diminished with decreasing force contributions of one leg pair.

### ii) Shape consistent force reduction with mid-stance as reference point

In general, the results are similar to those where only force amplitude was reduced. Some differences occur with regard to the height of the total force amplitudes. While at low duty factors, maximum force amplitudes are still about the same for all numbers of pairs of legs, amplitude excesses occur when force contributions of one leg were reduced; particularly between 2 and 1 pair of walking legs and high duty factors, i.e. relatively low speeds (Fig. 3). Due to these excesses, the slopes from intermediate phase values towards θ = 0.5 of the total force amplitude increase.

**Fig. 3.**
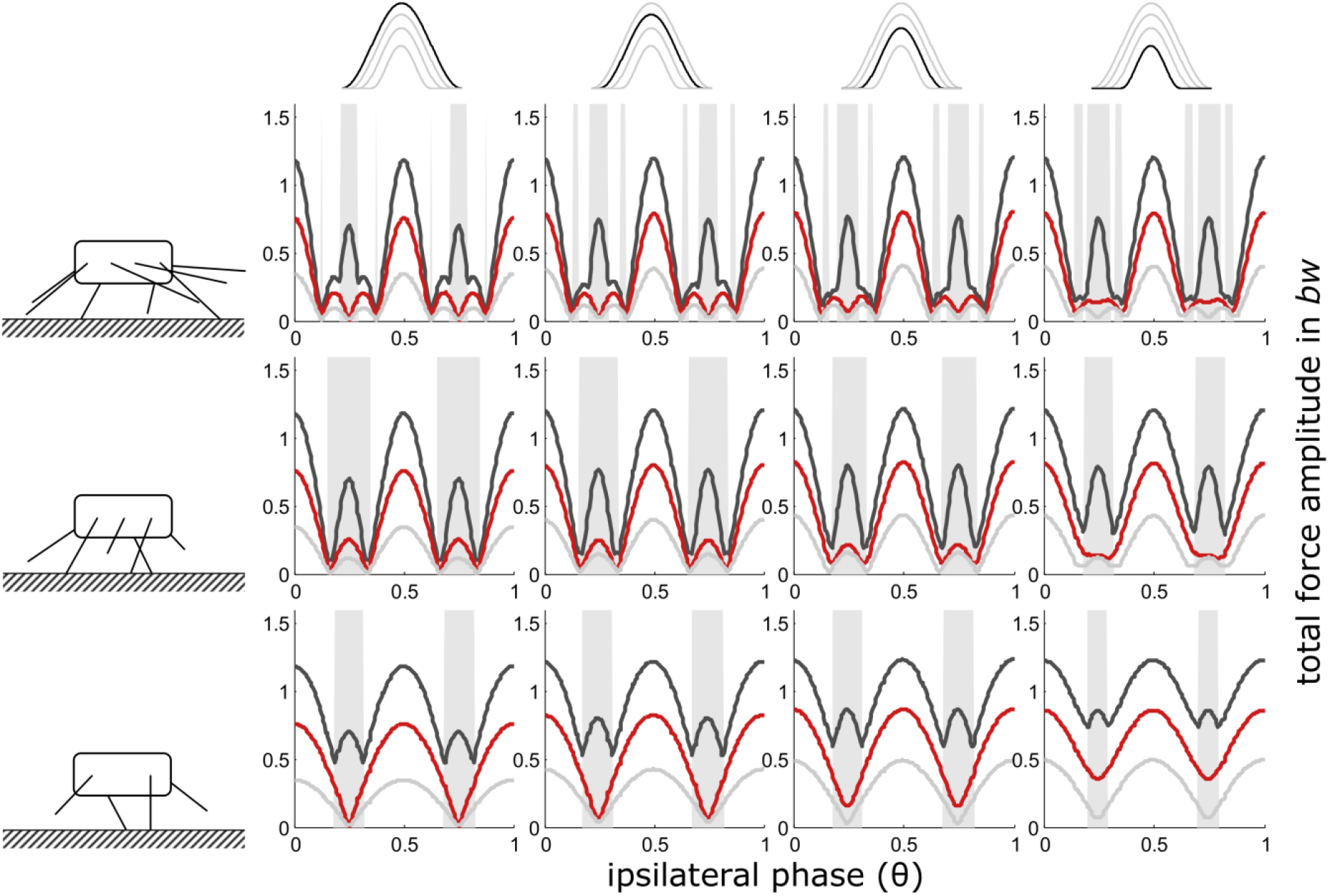
Mid-stance as reference point (iii): Dependency of total vertical leg force amplitudes on the phase shift of ipsilateral adjacent legs and duty factor with gradually decreasing contributions of increasing numbers of leg pairs. GRF decreased while their shape (ratio of width to height) remained constant i.e. also contact durations decreased with the reduction of the impulses beneath leg pairs. Duty factors shown: 0.3 (dark grey), 0.5 (red) and 0.8 (light grey). The top row indicates the degree of reduction in the pair of legs with reduced GRF and the type of force alignment (mid-stance (iii)). Accordingly, in the third column from the right, the impulse beneath one leg pair is reduced by ¼, in the 2^nd^ column from the right by ½ and in the rightmost column by ¾. The pictograms to the left indicate the number of propulsive pairs of legs valid for the respective row of subplots, i.e. four in the upper row, three in the middle row and two in the bottom row. With low duty factors and intermediate phase shifts, the peak frequency of the force oscillations assumed values higher than two times the stride frequency; these intervals are shaded in grey.

### iii) Shape consistent force reduction with take-off as reference point

As was generally the case with reduced contact durations, decreasing contact duration of one pair led to excesses of the total force amplitudes, particularly at higher duty factors and also when take-off or touch-down served as reference point for the timing of the legs with reduced GRF. However, in contrast to the previous condition, the occurrence of the total ground force maxima in the phase-range shifted away from θ = 0.5 towards higher phase values when reduced GRFs were aligned with the take-off of regular force traces (Fig. 4), and towards lower phase values when reduced GRFs were aligned with the touch-down (regime ii) of regular force traces (fig. S1). This shift was particularly pronounced between two and one pair of legs and high duty factors. At high speeds and low duty factors, the shift was almost negligible.

**Fig. 4.**
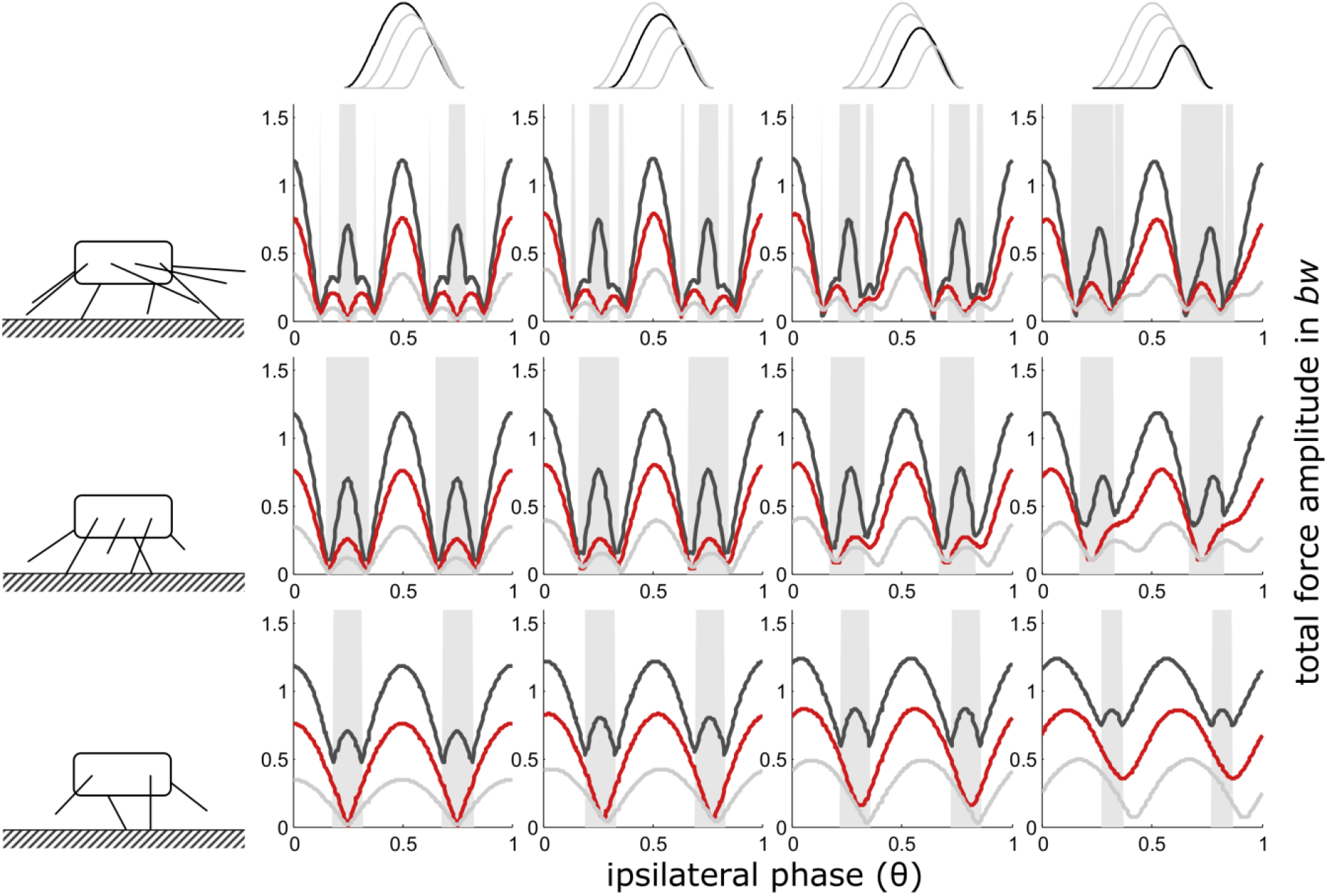
Take-off as reference point (iv): Dependency of total vertical leg force amplitudes on the phase shift of ipsilateral adjacent legs and duty factor with gradually decreasing contributions of increasing numbers of leg pairs. GRF decreased while their shape (ratio of width to height) remained constant i.e. also contact durations decreased with the reduction of the impulses beneath leg pairs. Duty factors shown: 0.3 (dark grey), 0.5 (red) and 0.8 (light grey). The top row indicates the degree of reduction in the pair of legs with reduced GRF and the type of force alignment (take-off (iv)). Accordingly, in the third column from the right, the impulse beneath one leg pair is reduced by ¼, in the 2^nd^ column from the right by ½ and in the rightmost column by ¾. The pictograms to the left indicate the number of propulsive pairs of legs valid for the respective row of subplots, i.e. four in the upper row, three in the middle row and two in the bottom row. With low duty factors and intermediate phase shifts, the peak frequency of the force oscillations assumed values higher than two times the stride frequency; these intervals are shaded in grey.

## Discussion

Figuring out the advantages and disadvantages of using different numbers of legs and why in some clades the number of propulsive legs can vary is just as challenging as finding out ultimate reasons for speed dependent changes in the number of propulsive legs as found in a range of multilegged species (cp. *17, 19, 25, 42*). In fact, the two questions seem to be intertwined. The present model helps to reveal some of the underlying mechanisms.

Though there are quite a few studies on the emergence of bipedalism (e.g. *43, 44, 45*), particularly in view of our own preferred mode of locomotion, changes in the numbers of propulsive legs are much less studied with regard to multi-legged animals. Nevertheless, the underlying mechanisms have tremendous importance for, both, understanding the drivers of evolutionary processes affecting the locomotor system of multi-legged organisms, as well as the meaningful development of swiftly moving robots with polypedal designs.

The present study shows that, in principle, the mechanical effects of changes in leg numbers follow a continuum (Figs. 2-4). Nevertheless, in quadrupedal locomotor systems, the reduction of GRF under some feet has decisively different consequences than similar reductions in systems with more than two pairs of legs. Therefore, the step from quadrupedal to bipedal locomotion seems to take place under somewhat different constraints.

Ancestral legged locomotor systems, just after a certain clade’s colonisation of land, were rather slow and did not exploit energy cycling (*21, 46*). With such conditions, ipsilateral phase relations deviating from 0.5, i.e. metachronal wave coordination, were an effective way to avoid energetically costly vertical oscillations that might even disturb vision and position control (*47-49*). In later developmental stages of terrestrial ecosystems, with established complex food-webs and accompanied coevolution towards efficient locomotion, synchronised sets of alternating legs became important means for energetic optimisation of locomotion and dynamic stabilisation in the sagittal plane (*8, 29, 41*).

With small numbers of legs, the alternating application of ground forces by sets of synchronised legs results in vertical oscillations of an animal’s body of two times stride frequency (*1*). These oscillations can be used for temporal storage of movement energy during the sets’ stance phases (see introduction). With temporally desynchronised sets of legs, vertical oscillations can be largely avoided (*3*). Accordingly, particularly in few-legged designs, synchronisation or desynchronisation of the legs in functional sets enables the choice between pronounced or small body oscillations, i.e. conditions that correspond best with the constructional constraints and mechanical requirements of a locomotor apparatus as well as particular environmental requirements (*37, 50-52*), both of which contribute to energetic optimisation.

Similar to the assumed evolutionary paths of legged animals, many extant species do not suddenly change their number of propulsive leg pairs, but instead do so gradually. Just as assumed, this gradual decline of the GRFs beneath some legs and the way in which it is achieved has some salient effects on the overall GRF and body dynamics.

It has been shown earlier, that the width of the central region in the phase graphs increases with decreasing numbers of leg pairs. This means that animals with a higher number of legs are less able to use vertical oscillations of the body for the energetic optimisation of their locomotion (*3*). However, in contrast to the general trend, at the transition from two to one propulsive leg pair, the position of the amplitude minimum in the phase space remains unchanged. Nonetheless, with gradually decreasing GRF under a pair of legs, the depth of the minima diminishes and the slope of the amplitude decrease continues to decline, particularly when contact durations of all legs were kept constant. Therefore, reducing GRF under one of the leg pairs increases the ability of quadrupeds to exploit vertical oscillations for energy recovery, even at phase relations where animals with more legs and even quadrupeds that equally distribute GRFs among their legs are affected by oscillation minima. However, this also means that oscillation minimisation can no longer be used as an energy optimisation tool. Lizards and cockroaches can serve as excellent examples to illustrate the effects. The American cockroach (*P. americana*) gradually reduces the number of propulsive leg pairs from three to one when speeding up to their maximum running velocity of about 1.5 ms^-1^ (*17*). At high running speeds, the legs reach cycle frequencies of up to 25 s^-1^ and faster runs are more likely affected by internal and external disturbances. Controlling such fast movements by reflex loops would require neuronal conduction speeds close to and beyond the physiological limits of the nervous system (*53-56*).

According to the predictions of the present model, the reduction of the GRFs under the forelegs and the gradual reduction of the number of leg pairs in contact with the substrate allows *P. americana* to increase the tolerance of its locomotor system against deviations from the ideal alternating leg coordination pattern, while still being able to effectively use vertical body dynamics for energy cycling. Interestingly, elasticities, allowing for efficient energy cycling, have been found precisely in the hips of the cockroaches’ rear legs (*57*), i.e. the pair of legs that propel the animals at maximum running speeds.

Locomotor systems with fewer legs tend to have more pronounced vertical oscillations (*8*). These oscillations, however, have been shown to be key for mechanically implemented stability mechanisms, i.e. self-stability (*58, 59*). The locomotor system of hexapedally running cockroaches does not seem to be dynamically stabilised in the sagittal plane (*60*). However, the gradually declining GRF of some legs increases the stability of vertical COM oscillations and therefore also the cycling of kinetic and potential energies in the sagittal plane. Accordingly, gradual reduction of propulsive legs seems to be a suitable way to increase the animals’ running stability against larger disturbances at high speeds. The same mechanisms seem to be at work in facultative bipedal lizards (cp. *47*). In these fast sprinters, the range of ipsilateral phase relations in which leg elasticities can be effectively exploited and the robustness against coordinative deviations increases significantly when the GRF of the forelegs diminishes. In quadrupeds, when changing from two to only one propulsive leg pair, the increased robustness against disturbances could possibly be the main reason for bipedalism at high running speeds.

Long hindlegs, which are typical for bipedal species, allow for longer contact phases and therefore increased power generation and higher maximum running speeds (*2*). Long hindlegs and backward shift of the COM often have co-evolved and have mutually conditioned each other (*61*). Correspondingly, tailless lizards cannot employ bipedal locomotion even if they would otherwise have been able to do so (*62*). However, originally tailless species, like some monkeys, great apes or technical implementations like humanoid robots, can bring their COM over the hind limbs by raising the body. Like in cockroaches, this automatically reduces the GRFs generated by the forefeet and increases the stability of vertical body oscillations (see above). However, increasing bipedalism decreases the ability of a locomotor system, be it biological or mechanical, to employ asymmetrical gaits, i.e. gallop gaits and the employment of the lumbar spine for functional elongation of the legs (*2*). This functional lengthening gives the galloping gaits an advantage over symmetrical gaits when it comes to maximum speed performance. Accordingly, quadrupeds must choose between increased inherent stability of bipedal locomotion and higher maximum speed of quadrupedal gallop like locomotion.

Lizards and cockroaches are also good examples exhibiting two different ways of reducing the forces under their feet. Thus, most animals that facultatively reduce the number of propulsive legs, do so either with the body axis basically in parallel with the substrate or with a more erect body position. In both cases, a gradual increase of body height or of the position of the foreleg hips respectively results in gradual loss of ground contact in the relatively shorter anterior legs. The GRFs of these legs diminish as their contact phases reduce and load is transferred to other legs. With increasing speed, in American cockroaches, the body’s angle of attack increases from about 0° to about 30° (*17*). While no phase shifts of the forelegs’ take-offs have been found (*63*), this results in reduced contact phases by increasingly delayed touch-downs and finally complete omissions of the ground contacts.

A range of lizard species that become bipedal at higher running speeds (e.g. *18, 42, 62*), keep their body axis largely constant in parallel with the substrate (*18*), which seems to be necessary in order to stabilise the vertical position of the head, supporting vision and vestibular perception (*47*). When speeding up, they gradually increase the height of their body above the substrate and transfer body weight onto the longer hind limbs (*18, 64*)(https://www.youtube.com/watch?v=ExyMxKDxT9M). This results in diminishing contact durations of the forelegs, but in contrast to the cockroach, here, the timing of the force maxima does not necessarily shift. A similar pattern of force reduction can be assumed for likewise facultative quadrupedal dinosaurs of the hadrosaurid, iguanodontid or ceratopsid clades.

Since the model is principally insensitive against the position of the leg pair under which GRF is reduced, shortened stance phase durations due to early termination may find its equivalent in more rearward directed legs. Thus, in fast running spiders of the species *Ancylometes bogotensis* the contact durations of the forelegs (about 79%) as well as that of the rear legs (about 83%) are significantly shorter than that of the third and second legs (*26*).

However, exclusive amplitude reduction while contact durations are maintained might also be relevant in the locomotion of animals that do not necessarily reduce the number of propulsive leg pairs, particularly when anatomical characteristics cause an unbalanced weight distribution between rear and forelegs. Such imbalances are known for a number of ungulates, where typically the forelegs bear the greater weight (e.g. *65, 66*), but also for crocodiles, where the heavy tails cause higher load on the rear legs (*67*). Roughly equal contact durations for all propulsive legs, then, would facilitate a constant position of total force amplitude maxima at θ = 0.5 (Fig. 2) and therefore predictable responses of total GRF and body dynamics to ipsilateral phase changes. The same applies to robots with uneven weight distribution either being caused by the mass distribution within their body or when carrying loads (see introduction).

Contrary to this, shorter contact phases of some legs result in more slender total force trajectories when the force maxima of all legs of a set coincide. This is particularly pronounced at the transition from two to one pair of legs, since with low numbers of leg pairs the characteristics of a single force trace have greater impact on the total GRF. Accordingly, emerging total force excesses are more pronounced in locomotor systems with fewer legs (fig. S3). Moreover, additional phase shifts induced by shortening contact durations (*θ*_*add*_, see methods) occupy larger proportions of a stride when duty factors are high, which results in pronounced total force excesses at medium to high duty factors. Since under these conditions the temporal overlap of the consecutive sets of legs is maximized and absolute COM oscillations are low, this should rarely impact running dynamics negatively.

With reference points other than the position of the force maximum (i.e. touch-down (ii) or take-off (iv)), decreasing contact durations of single leg pairs also led to shifts in the occurrence of total force amplitude maxima and minima. Also here applies, the lower the number of pairs of legs involved in the force interference and the higher the duty factor, the larger are the phase shifts of the amplitude maxima and minima. With more than two pairs of legs, this shift can also lead to asymmetries of the central region of the phase diagram were the force amplitude decline on one side is steeper than the other. These slope differences might be the reason for differences in the direction of phase deviations as observed in previous studies. Thus, when blaberid cockroaches change from alternating to high-speed metachronal leg coordination, they shift ipsilateral phases towards lower phase values (*32*), while mites shift towards higher values (*1*). When islandic horses use their notorious tölt gait, they also shift towards lower phase ratios (*4*) with their forelegs bearing higher loads than their rear legs (*68*). It seems conceivable that phase shifts are preferably adapted in the direction that enables smaller deviations from the alternating coordination pattern and thus reduces vertical oscillations more efficiently. Unfortunately, the lack of GRF measurements in blaberid cockroaches running at higher speeds with distinct metachronal leg coordination and even more so in mites, prevents further comparative inferences.

However, by reversing the reasoning, animals and legged machines aiming at smooth COM trajectories by using ipsilateral phase relations deviating from θ = 0.5 can experience sudden force and COM amplitude increases when the contact durations of some legs are accidentally delayed or terminated prematurely. Shortened contact durations can cause a sudden shift of the maximum total force amplitude away from the alternating coordination pattern, which has the potential to destabilise locomotion dynamics. Fortunately, pronounced phase shifts and therefore high relative changes in total force amplitudes are to be expected only at higher duty factors and lower speeds, where COM amplitudes are generally small. At higher speeds, the position of the total force amplitude maxima deviates only little from θ = 0.5. Interestingly, gradual reductions in the number of propulsive legs occur mainly when animals shift to higher running speeds (*17, 19, 42*), i.e. when locomotor systems are less vulnerable against truncated contact phases.

Emerging force excesses (see fig. S3-4) are also a result of the way normalisation is implemented in the model. As the summed-up forces are divided by their mean value (see methods), an increased slenderness of the total force trajectory results in force amplitudes exceeding the values found for integer numbers of pairs of legs and unchanged contact durations. Nevertheless, in real animals and legged machines, total GRFs have to be strong enough to counteract gravitational forces, such that, under natural conditions, more slender total force traces would also lead to force maxima that exceed those of stouter ones.

At intermediate phase relations, i.e. when total force amplitudes are usually low, total forces and body dynamics accordingly are also affected by oscillation frequencies higher than the usual two times stride frequency (Figs. 2-4). Particularly at low duty factors, these oscillations can have significant amplitudes. Unfortunately, their energy content cannot be reconverted into forward movements (*69*). However, frequencies higher than twice the stride frequency are usually effectively dissipated due to the damping properties of the leg muscles and other internal structures (*3, 21, 57, 70, 71*), which is why these high frequencies are practically unobservable in animals. Nevertheless, in underdamped machines it can become a problem and should be taken into account.

Although the broad central regions of the amplitude-phase dependency in quadrupeds enables significant deviations from the ideal alternating coordination pattern without losing the ability to recover movement energy in the late stance phase (see above), the occurrence of harmonics, however, has the potential to hamper efficient locomotor dynamics. Nonetheless, particularly with strongly reduced GRF under one leg pair, the extent of high frequency oscillations decreases (Fig. 2-4), making it tolerable to slip into the range were harmonics normally occur.

Facultatively bipedal species such as a range of lizard species (*61*), monkeys (*72*), bipedal species like birds (*73, 74*) but also polypedal animals like ants (*75*) or spiders (*26*) exhibit single leg GRFs deviating from the symmetrical cosine shapes used in the present approach. These deviations are required either to counteract posture induced pitch moments or they are caused, in polypedal species, by interacting horizontal leg force components as proposed by the lateral bracing hypothesis suggested for arthropods moving with alternating sets of synchronized legs (*21*). In particularly slow moving species like tortoises, skewed vertical ground forces can be caused also by the contraction properties of the driving muscles (*29*). However, when single leg GRFs have tails to one side, either prior to or after the vertical force maximum, these tail forces are small if compared to the force maximum. Accordingly, they are probably of secondary importance for the force interference pattern, total body oscillation frequencies and amplitude but certainly deserve further examination since such deviating GRF shapes may affect the timing of the force maximum. Although timing effects are already mapped by the present approach (paradigms ii - iv), here only temporal deviations in the force-reduced legs are considered whereas temporal shifts of the force maximum in the remaining legs can result in additional changes in the phase dependency of the total force amplitudes. Otherwise deviating GRF shapes such as the two-humped patterns found in walking bi- and quadrupedal animals (*69, 76*) might lead to higher than expected oscillation frequencies, i.e. they would appear as harmonics in the present approach. However, although central dips can, in principle, compromise the determination of the dominant vertical oscillation frequency, these dips are naturally smaller than the vertical forces of the single legs themselves, i.e. smaller than the main signal. Accordingly, deviations from the results of the present approach are likely to be small.

## Conclusion

The reductionist approach and abstraction from the huge manifold of anatomical and control solutions of biological and technical legged locomotor apparatuses enables an overarching view. For arbitrary numbers of walking legs and all possible ipsilateral phase relations the model provides total force amplitudes and oscillation frequencies resulting from leg force interference. Real animals do usually use only small fractions of the generally possible combinations, which impedes mapping the entire solution space.

Despite its simplicity, the model provides insight in the effects of gradual ground force reduction. The results show that the transition from quadrupedal to bipedal designs differs from those between different polypedal designs. In quadrupeds only, the gradual force reduction in one pair of walking legs stabilises the vertical oscillations of total vertical GRF, which is a prerequisite for dynamic stabilisation of locomotion in the sagittal plane. Moreover, for certain conditions (paradigms ii and iv) the model shows, that total force maxima and minima can shift away from the positions they occupy with all legs exerting the full amount of GRF. Since these extremums’ positions within the phase space appears to be causative for the occurrence of certain symmetrical gaits in a number of animals (*3*), the timing of legs with reduced GRF may provide a new perspective to help understand how and why specifically constructed animals adjust their leg coordination patterns.

## Methods

The present phenomenological model approach is based on vertical forces exerted by individual legs on the ground (*3*). The leg force interference approach, on which the present model is based, examines the amplitude of the total vertical force component and how it changes with changing ipsilateral phase relations (from 0 to 1). Here this examination is carried out for three different duty factors (0.3, 0.5 and 0.7). With decreasing duty factors total GRF amplitudes increase, except at such ipsilateral phases were leg force interference leads to diminishing force amplitudes (cp. Fig. 1), which is due to shorter stance phases that require higher GRF to balance body weight.

By following Weihmann (*3*), when calculating interferences of the single legs’ vertical GRFs, all forces were summed up for an interval of 20 strides of the rear legs and divided by the mean of this sum, which resulted in total force oscillations normalised onto body weight. That is, the GRFs of all legs taken together just counterbalance the gravitational forces acting upon the animal’s body. Afterwards, the resulting force oscillations were subjected to spectral analyses, which were accomplished by using a Fast Fourier Transformation algorithm. All analyses were performed using MatLab scripts (MATLAB 7.10.0; The MathWorks, Natick, MA, USA). Resulting amplitudes correspond to ½ of the peak to peak values of the total ground forces, i.e. the difference between the absolute maximum and minimum.

Initially, all GRFs have equal amplitudes for all walking legs and are applied for the same time intervals. Such patterns of force application and temporal distribution are supported by a wide range of experimental findings (*29*) as well as model approaches focussed on spring mass dynamics (*77*). According to these studies, force traces were assumed as symmetrical, i.e. leg loading and unloading required the same interval of time. In general, the forces applied during the legs’ stance phases were modelled as 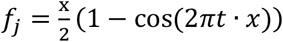 for the time interval t from 0 to take off, i.e. the start and the end of the stance phase (cp. Fig. 1, fig. S2); with x being 1 in regular steps and when impulse reduction was reached by reduction of force amplitude alone. The number of a pair of legs is indicated by the index *j*.

In contrast to previous work, here the GRFs of the respective anterior legs, i.e. those with the lowest index, were gradually reduced in defined steps of ¼ of the initial impulse, i.e. the area under the force trace. Hence, after four reduction steps, this resulted in a reduction of the number of leg pairs by one. Starting with four pairs of legs, the reduction continued until there was only one pair left. In real locomotor systems, the reduction would require a transfer of body weight to the remaining legs. Thus, force reduction in one pair of legs would result in increased GRFs in the remaining legs. However, with shape maintenance of the GRFs (see below) a force transfer leads to equivalent results if the differences between the impulses of the force reduced and the remaining legs were kept constant at multiples of ¼ and the time axis is adjusted such that despite effectively increasing stride, contact and swing duration, the stride lengths were set to 1. The resulting geometrically similar single leg force traces interfere equivalently and due to the normalization onto body weight, the resulting total force trajectories provide equivalent frequency spectrums and amplitude-phase dependencies.

With constant contact durations, the transfer of force towards the remaining legs results in increasingly steep single leg ground forces, i.e. an additional shape distortion of the GRFs affecting the interactions between the leg forces prior to normalization onto body weight. This leads to strong total force excesses when the legs are synchronised. However, additional degrees of freedom like additional shape distortions should be avoided here which is why stride duration and force amplitude of unaffected legs was generally set to 1, which is equivalent to a previous approach with integer numbers of leg pairs (*3*). Concurrently, this approach facilitates comparability between locomotor apparatuses with different numbers of leg pairs and the comparison with experimentally obtained data.

Phase dependent changes in the total force amplitudes and resultant vertical body oscillations are unlikely to be useful with leg numbers above 8, i.e. four pairs of legs (*3*) and the model is not applicable if no ipsilateral phase relations can be applied, i.e. with only one pair of legs left. Therefore, Figs. 2-4 and S3 show only the range from just above one active pair of legs to four pairs of legs. Nevertheless, the model applies to arbitrary numbers of leg pairs, as illustrated in fig. S4 (see supporting information).

When contact durations were kept constant and impulse reduction was achieved by reducing force amplitudes alone, x was unaltered and the force amplitudes beneath the forelegs reduced stepwise by ¼ of the initial value by multiplication with 0.75, 0.5, 0.25 or 0, respectively. When the general shape of the force trace (i.e. width to height ratio) was maintained, the values for x assumed the square root of these values, specifically 0.866; 0.7071 and 0.50, in the pair of legs with reduced GRF. This resulted in shape consistent reduced impulses of ¾, ½ and ¼ of a regular step (fig. S2).

Strides are comprised of an initial stance and a subsequent swing phase where no forces are applied onto the ground. The quotient of contact duration and stride duration defines the duty factor β of a leg. Initially, stride duration, duty factors and ground forces were equal for all legs. In the present study, three different duty factors and their impact on total force and body dynamics are explored. The chosen values of 0.3, 0.5 and 0.7 are representative in a wide range of species for fast, medium and slow runs (*29*). Keeping the shape of the GRF traces constant and reducing the impulse of some legs affects stance and swing durations as well as the duty factor of the respective legs, and also the timing of touch-down, take-off or the occurrence of the force maximum (see below).

The phase relations between legs were generally defined with regard to standard stride durations, which were set to 1. The ipsilateral phase shift θ was determined by the occurrence of one leg’s touchdown within the stride period of an adjacent leg; and assumed values between 0 and 1. Owing to the circular nature of the distribution of phase relations, phase shifts of 0 and 1 are equivalent (*30*); each indicating the synchrony of the ipsilateral legs. However, due to static requirements and considerations, alternating sets of legs are more common in nature and technology.

In accordance with Weihmann (*3*), contralateral phase relations between the rear legs, i.e. those legs with the highest index, were set to 0.5, i.e. they alternated strictly, which, in quadrupedal animals, is a characteristic of symmetrical gaits (*4*). However, contralateral phase relations of about 0.5 are also often found in arthropods (*21*). In order to generate different leg coordination patterns, the θ of all ipsilateral legs were uniformly shifted from 0 to 1, which resulted in changing contralateral phase shifts in all leg pairs anterior to the rear legs. Whether the hind legs or the front legs served as pacemakers makes no difference. Although, in the model, force reduction was always started at leg pair 1 and proceeded to subsequent pairs of legs, the model is indifferent to the location of the force-reduced leg pair, since leg positions are not defined; therefore the model can consider reduced force in any leg. Ipsilateral phase, duty factor, number of leg pairs and x (i.e. the variable determining the impulse beneath the most anterior pair of legs), were the only changed parameters. Oscillations of total vertical forces, then, emerged from the interplay of these measures.

Reducing the impulse of legs while maintaining the shape of the force trace, resulted in shorter contact durations and sometimes led to additional temporal shifts. Since θ was defined according to touchdown and stride duration, kinematic phase was subject to additional phase shifts when shortened stance phases were aligned with mid-stance or take-off of the standard stance phases. Numerically, additional shifts were implemented prior to any regular ipsilateral phase shift as described above. If reduced stance durations were aligned with mid-stance, such that force maxima of the shortened and the standard stance phase concurred, the actual phase increased by an additional phase shift of 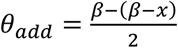. With alignment at take-off, the additional phase shift was *θ* = *β* − (*β* − *x*).

## Supplementary Material

**fig. S1.**
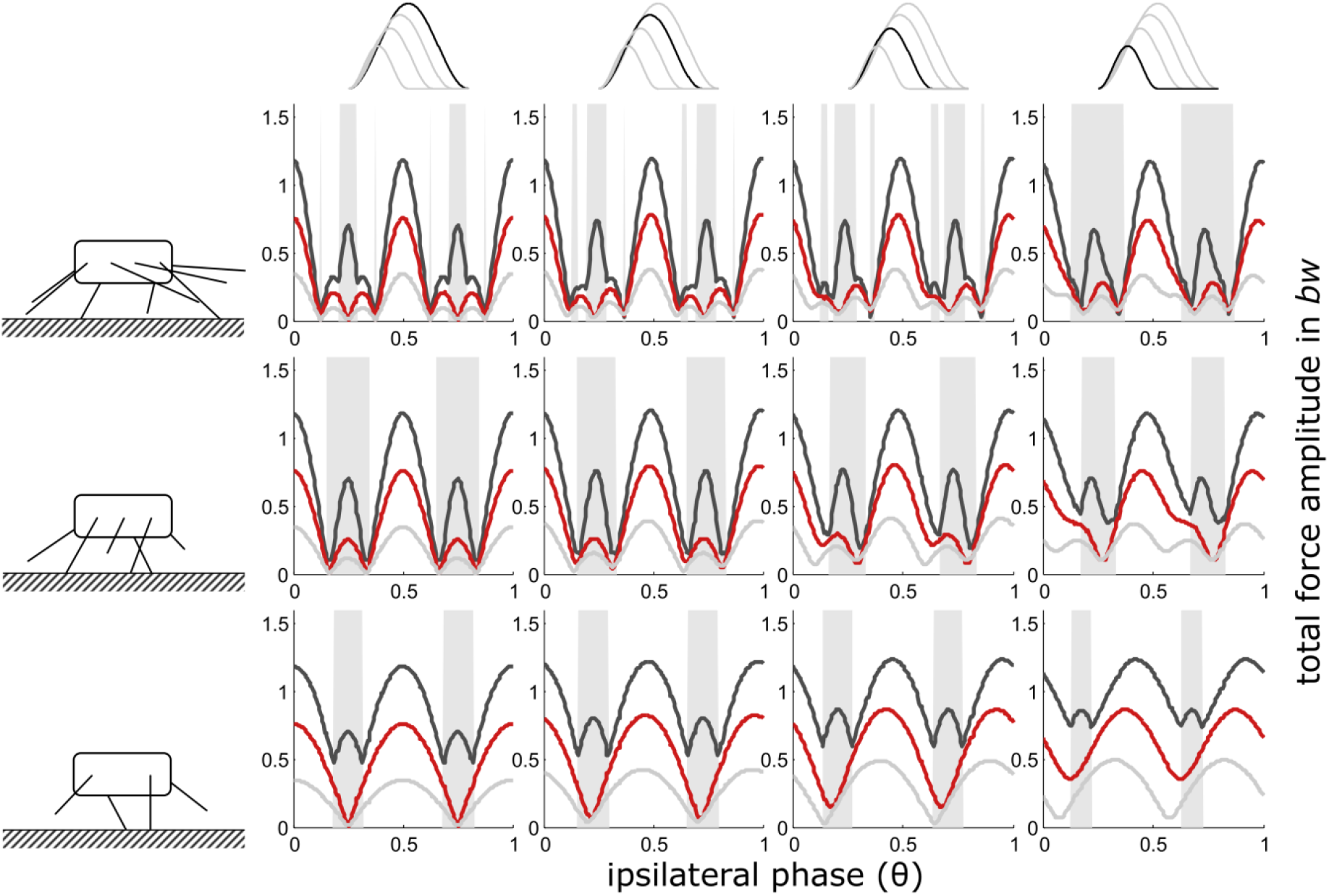
Touch-down as reference point (ii): Dependency of total vertical leg force amplitudes on the phase shift of ipsilateral adjacent legs and duty factor with gradually decreasing contributions of increasing numbers of leg pairs. GRF decreased while their shape (ratio of width to height) remained constant i.e. also contact durations decreased with the reduction of the impulses beneath leg pairs. Duty factors shown: 0.3 (dark grey), 0.5 (red) and 0.8 (light grey). The top row indicates the degree of reduction in the pair of legs with reduced GRF and the type of force alignment (touch-down (ii)). Accordingly, in the third column from the right, the impulse beneath one leg pair is reduced by ¼, in the 2^nd^ column from the right by ½ and in the rightmost column by ¾. The pictograms to the left indicate the number of propulsive pairs of legs valid for the respective row of subplots, i.e. four in the upper row, three in the middle row and two in the bottom row. With low duty factors and intermediate phase shifts, the peak frequency of the force oscillations assumed values higher than two times the stride frequency; these intervals are shaded in grey.

**Fig. S2.**
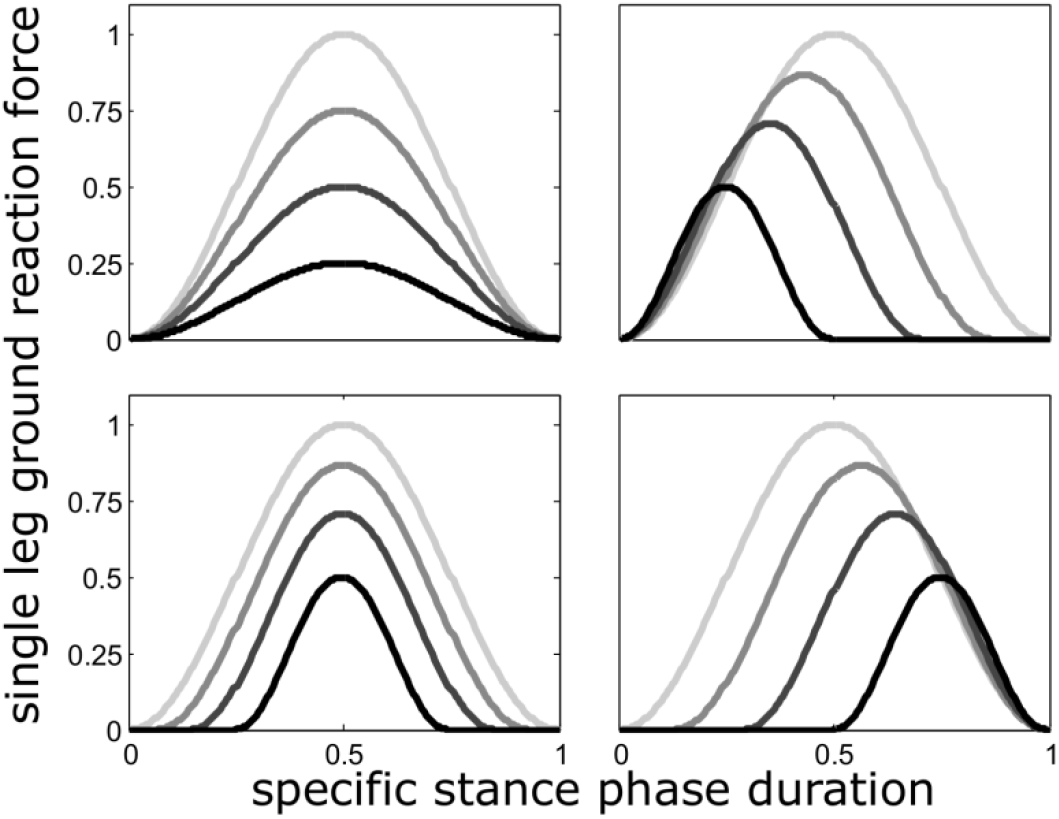
Four different ways in which GRF reductions were implemented. Every force reduction step results in an impulse reduction of ¼. GRF reduction was implemented either by decreasing force amplitudes only (upper left; scheme i) or by a combined reduction of contact duration and force amplitude (schemes ii to iv; see methods). When GRF reduction followed the latter, reduced forces were aligned in three different ways to the GRF of the remaining legs. Reduced GRF were aligned for θ = 0.5 either to touch-down (top right), mid-stance (bottom left) or take-off (bottom right) of the legs with normal GRF.

**fig. S3.**
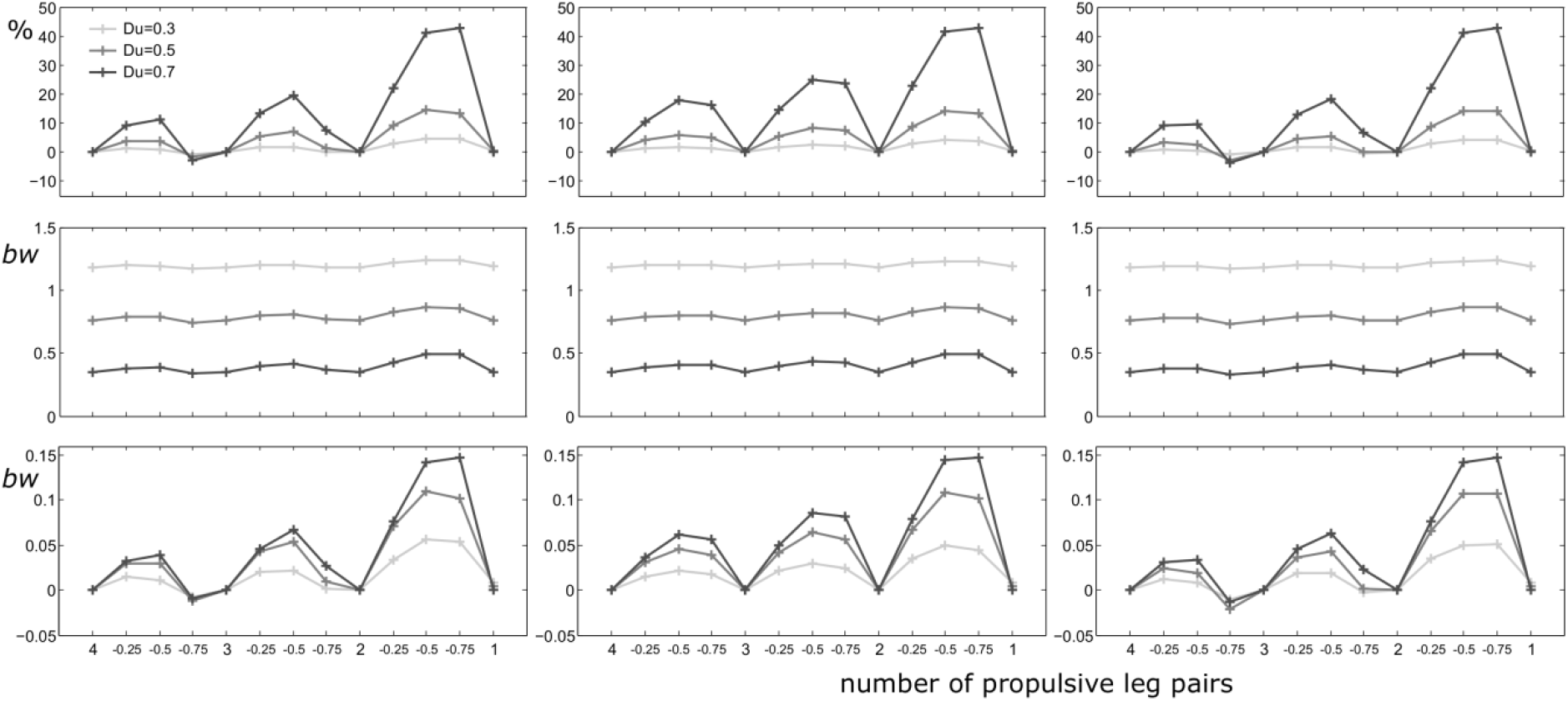
Total force excesses for shape consistent force reduction. Left: take-off alignment (iv); Middle: mid-stance alignment (iii); Right: touch-down alignment (ii). The abscissa gives the number of propulsive pairs of legs (large numbers) and the degree of force reduction (small numbers). Results are shown for duty factors of 0.3, 0.5 and 0.7 (see legend). Upper row: relative force excess to integer numbers of leg pairs in %; second row: absolute maxima of the force amplitude values (at θ = 0, 0.5 or 1); bottom row: force difference in body weight.

**fig. S4.**
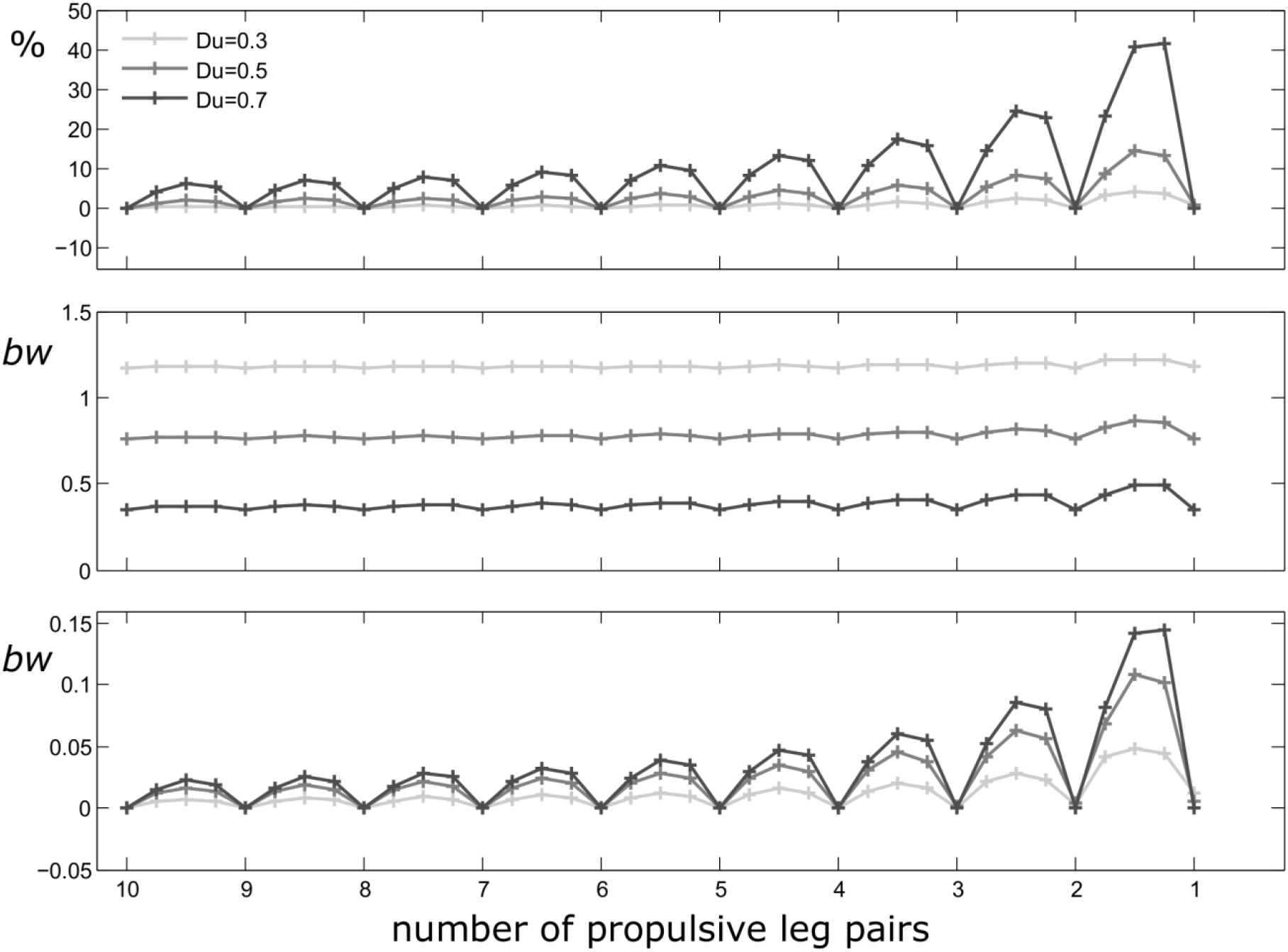
Total force excesses for shape consistent force reduction and mid-stance alignment (scheme iii), for continuous GRF reductions from a locomotor apparatus with 10 pairs of legs down to only one pair of legs. The abscissa gives the number of propulsive pairs of legs. Results are shown for duty factors of 0.3, 0.5 and 0.7 (see legend). Upper row: relative force excess to integer numbers of leg pairs in %; second row: absolute maxima of the force amplitude values (at θ = 0, 0.5 or 1); bottom row: force difference in body weight.

## Data sharing plans (for all data, documentation, and code used in analysis)

All data are contained in the manuscript. Code and scripts will be provided upon request.

